# Prenatal Cannabinoids Produce Sex-Specific Changes in Risk Assessment and Shared Increases in Repetitive Behavior in Adult Offspring

**DOI:** 10.1101/2025.10.23.682511

**Authors:** Alba Cáceres-Rodríguez, Daniela Iezzi, Pascale Chavis, Olivier J. Manzoni

## Abstract

The widespread perception of cannabinoids as harmless remedies has led to increasing use of both cannabidiol (CBD) and Δ9-tetrahydrocannabinol (THC) during pregnancy, yet their long-term impact on offspring behavior remains incompletely characterized. To directly compare these compounds, pregnant mice received daily CBD or THC (3 mg/kg, gestational days 5–18) and adult progeny of both sexes were evaluated (P90–120). To capture complementary dimensions of emotional and defensive responding, we employed the elevated plus maze (EPM) to quantify anxiety-related measures and ethological risk-assessment postures (e.g., stretch-attend) and the marble burying (MB) test to index repetitive/defensive behavior; together these assays provide convergent but distinct readouts that increase sensitivity to cannabinoid-induced effects. In the EPM, prenatal CBD exposure selectively increased stretch-attend postures in females, indicating heightened risk assessment; males were unaffected, and THC produced no comparable effect. Classical EPM indices—time in open or closed arms, preference for open-arms, and total distance traveled—were unchanged across groups, except that THC-exposed females exhibited hyperlocomotion. In contrast, marble burying was increased in both sexes following prenatal exposure to either CBD or THC, indicating a shared enhancement of repetitive/defensive responses. These results show that prenatal cannabinoid exposure yields enduring, sex-dependent alterations in adult behavior, with CBD selectively heightening female risk assessment and both cannabinoids increasing repetitive behavior across sexes, challenging the notion that CBD is a benign alternative to THC during gestation.

## Introduction

Many psychiatric and neurodevelopmental disorders that emerge in adolescence or adulthood— including anxiety, depression, bipolar disorder, schizophrenia, and obsessive–compulsive disorder—are now understood to originate in fetal brain development, a period of heightened vulnerability to environmental disruption. Prenatal exposure to psychoactive substances such as cannabis is of particular concern, as both Δ9-tetrahydrocannabinol (THC) and cannabidiol (CBD) readily cross the placenta and can interfere with the maturation of neural circuits implicated in these conditions (Scheyer et al. 2019; Hurd et al. 2019; Johnson et al. 2022; Baker et al. 2018).

Cannabis is the most widely used illicit psychoactive substance among pregnant women, with prevalence rates of 5–10% in North America and even higher in regions where recreational use is legal (Young-Wolff et al. 2019; 2019; Volkow et al. 2019). Reported motivations include relief from nausea and vomiting, sleep disturbances, anxiety, pain, and mood symptoms (Brown et al. 2017). Both THC and CBD cross the placenta (Eidelman 2023) and are detectable in breast milk (Baker et al., 2018), thereby exposing the fetus and neonate during critical windows of brain development. While THC is primarily consumed for its psychoactive and antiemetic properties, CBD is increasingly marketed as a “natural,” non-intoxicating alternative and is often perceived as safer for use during pregnancy (Crume et al. 2018). Recent surveys indicate that up to 20% of pregnant women in the United States and Canada report using CBD-only products (Hammond et al. 2022), underscoring the urgency of evaluating its developmental impact.

THC exposure during neurodevelopment can disrupt the endocannabinoid system, which plays a central role in brain formation and maturation (Harkany et al. 2007), thereby predisposing offspring to psychiatric vulnerability (Hurd et al. 2019; Scheyer et al. 2019). Human studies support this view: gestational THC exposure has been associated with lower mental development scores in infancy (9 months), impaired verbal reasoning and short-term memory in early childhood (3–4 years), and, in females specifically, increased inattention and aggressivity at 18 months. Beyond infancy, prenatal cannabis exposure has been linked to heightened aggression, rule-breaking behaviors, and an elevated risk of psychotic-like experiences by age 10. Moreover, gestational cannabis exposure has been implicated in increased sensitivity to drugs of abuse later in life, potentially compounding pre-existing psychiatric risk (Manduca and Trezza 2022).

Rodent models provide converging evidence. Prenatal THC alters early developmental milestones, disrupts endocannabinoid signaling, and induces long-lasting changes in brain structure and function. During adolescence, exposed offspring exhibit deficits in social interaction, recognition memory, and emotional regulation, accompanied by molecular and electrophysiological alterations in corticolimbic circuits (Scheyer et al. 2019; 2023). Mechanistically, these outcomes are linked to disrupted glutamate and dopamine signaling in the mesocorticolimbic system and impaired cortical synaptic plasticity (Bara et al. 2018; Scheyer et al. 2019; Hurd et al. 2019; Hurd 2020). Effects are sex-dependent: loss of endocannabinoid LTD following prenatal THC exposure is observed only in males (Bara et al. 2018), whereas lactational THC exposure renders both sexes sensitive (Scheyer, Borsoi, Wager-Miller, et al. 2020; Scheyer, Borsoi, Alicot, et al. 2020, Scheyer 2019). In adulthood, prenatal THC exposure has been associated with persistent anxiety-like behavior, deficits in cognitive flexibility, and altered reward sensitivity, again with evidence for sex-specific vulnerabilities (Luján et al. 2024; Sarikahya et al. 2023; Frau et al. 2019).

Although CBD lacks the overt intoxicating effects of THC, preclinical studies indicate that prenatal CBD exposure also produces enduring neurobehavioral alterations. Rodent models report changes in anxiety-related behavior, memory, sensory gating, and pain sensitivity, often in a sex-dependent manner (Swenson et al. 2023). Extended exposure spanning gestation and early postnatal life alters repetitive and hedonic behaviors in adulthood (de Salas-Quiroga et al. 2020). Our own prior work demonstrated that prenatal CBD exposure can affect early growth trajectories, communication, and discrimination skills, as well as later electrophysiological activity in cortical regions involved in sensory and emotional processing, establishing a sex-specific trajectory suggestive of heightened anxiety in adulthood (Iezzi, Cáceres-Rodríguez, Pereira-Silva, et al. 2024; Iezzi et al. 2022; Iezzi, Cáceres-Rodríguez, Chavis, et al. 2025).

The present study directly compares the long-term behavioral consequences of prenatal THC and CBD exposure in male and female mice. Using two well-established assays—the elevated plus maze and marble burying—together with detailed ethological analysis of stretch-attend posture, we assessed anxiety-related, repetitive, and risk-assessment behaviors in adulthood. Our results reveal sex-specific vulnerabilities to CBD, with female offspring showing increased risk-assessment behavior, and a shared increase in repetitive behaviors across both sexes following prenatal exposure to either THC or CBD. These findings provide new insight into the distinct and overlapping consequences of prenatal THC and CBD exposure, with direct implications for public health messaging around cannabis use in pregnancy.

## Materials and methods

### Animals

All procedures complied with the European Communities Council Directive (86/609/EEC) and the United States NIH Guide for the Care and Use of Laboratory Animals and were approved by the French Ethical Committee (APAFIS #49376-2024051414491391). Adult male and female C57BL/6J mice (7-10 weeks old; Charles River) were housed in standard wire-topped Plexiglas cages (42 × 27 × 14 cm) under controlled conditions (21 ± 1 °C; 60 ± 10% relative humidity; 12 h light/dark cycle), with food and water available ad libitum.

After a one-week acclimation, pairs of females were introduced to a single male in the late afternoon. The presence of a vaginal plug was designated as gestational day 0 (GD0), after which pregnant females were housed individually. From GD5 to GD18, dams received daily subcutaneous injections of either vehicle, 3 mg/kg CBD (NIDA Drug Supply Program), or 3 mg/kg THC (Sigma-Aldrich). Cannabinoids were dissolved in a vehicle solution of Cremophor EL, ethanol, and saline (1:1:18) and administered at 4 mL/kg. Control dams (“Sham”) received the vehicle alone. The day of birth was designated postnatal day 0 (PND0). Pups were weaned at PND21 and housed separately by sex. Behavioral experiments were conducted in adulthood (PND90–120). Sample sizes (n) for each group are reported in the figure legends. Groups were formed from the complete offspring of 3–4 litters, as culling was not performed for ethical reasons.

### Behavioral Testing

To evaluate anxiety-related and repetitive behaviors in adulthood, we employed the EPM and the MB test (Iezzi, Cáceres-Rodríguez, Chavis, et al. 2025).

The elevated plus maze (EPM) consisted of two open arms (30 × 5 cm) and two closed arms (30 × 5 × 12 cm) extending from a central platform (5 × 5 cm). Each mouse was placed on the central platform and allowed to explore for 5 min. Sessions were recorded with a camera positioned above the apparatus, and behavior was analyzed using EthoVision software (Noldus, The Netherlands). Locomotor activity was quantified as total distance traveled (cm) and velocity (cm/s). Anxiety-related measures included time spent in the open arms, closed arms, and center, as well as relative open-arm preference, calculated as (time in open arms / total session time – time in open arms) × 100. Ethological parameters were assessed by scoring the proportion of time spent in normal, contracted, and stretched postures across the entire arena and within each maze subdivision (center, open arms, closed arms).

The marble burying (MB) test was conducted in a clean cage containing 4 cm of bedding, with 20 marbles evenly arranged on the surface. Each mouse was placed in a corner to initiate the trial, and behavior was observed for 20 min. At the end of the session, the number of marbles buried was recorded. A marble was considered buried when more than two-thirds of its surface was covered by bedding.

#### Statistical Analyses

were performed using GraphPad Prism 10. Data sets were first tested for normality with the D’Agostino–Pearson and Shapiro–Wilk tests, and potential outliers were identified using the ROUT method. Depending on the design, differences between medians were evaluated with two- or three-way ANOVA, followed by Sidak’s multiple-comparison post hoc tests as specified in the figure legends. Statistical significance was reported as exact p values in the figures. Experimental results are described qualitatively in the main text, while full statistical details— including sample size (N), test type, and p values—are provided in the figure legends. Quantitative data are presented as box-and-whisker plots showing median, minimum, and maximum values, with individual data points superimposed as scatter plots.

## Results

We employed the elevated plus maze (EPM) and marble burying (MB) to assess complementary facets of emotional and defensive behavior at adulthood. The EPM quantifies anxiety-related responses and ethological risk-assessment strategies such as stretch-attend postures, whereas the MB test probes repetitive and defensive actions associated with anxiety and compulsivity (Rodgers and Dalvi 1997; de Brouwer et al. 2019). Together, these paradigms provide convergent yet distinct behavioral readouts, enhancing sensitivity to cannabinoid-induced alterations and enabling us to distinguish overlapping from domain-specific effects of prenatal THC and CBD exposure.

### Absence of adult EPM open-arm effects after prenatal CBD or THC exposure

Epidemiological studies indicate that children exposed to cannabis in utero exhibit higher rates of anxiety-related problems later in life (Grant et al. 2018; Bolhuis et al. 2018; El Marroun et al. 2018). Preclinical work has similarly shown that gestational exposure to THC (Sarikahya et al. 2023) or to high doses of CBD (30 mg/kg) (Swenson et al. 2023) can increase anxiety-like behavior in rodents during adolescence and adulthood. In recent studies, we showed that in-utero exposure to a low dose of CBD produced sex-specific behavioral alterations in neonatal pups (Iezzi et al. 2022) and and anxiogenic-like phenotype in adolescent females (Iezzi, Cáceres-Rodríguez, Chavis, et al. 2025). Using the same prenatal exposure protocol, we asked whether the anxiogenic-like effects observed in adolescence persist into adulthood and extended the comparison to include prenatal THC.

In the EPM, a validated assay of anxiety-related approach–avoidance, no group differences were detected by sex or prenatal exposure for classical indices of anxiety: time in open arms, time in closed arms, and open-arm preference (Figure 1; Table 1 reporting all measured parameters). The only exception was a significant increase in total distance travelled by THC-exposed females, indicating that these offspring exhibited increases locomotor activity without accompanying changes in canonical anxiety-related measures. This absence of effect is consistent with our earlier report that prenatal THC does not alter open-arm exploration in adult rats of either sex (Bara et al. 2018). We recently showed that, under the same prenatal CBD regimen, female offspring exhibited an anxiogenic-like phenotype during adolescence, spending significantly less time in the open arms of the EPM (Iezzi, Cáceres-Rodríguez, Chavis, et al. 2025). In contrast to our recent report showing that, under the same prenatal CBD regimen, female offspring exhibited an anxiogenic-like phenotype during adolescence—spending significantly less time in the open arms of the EPM (Iezzi, Cáceres-Rodríguez, Chavis, et al. 2025)—these alterations were no longer evident in adulthood when assessed using standard EPM metrics.

**Figure 1.**
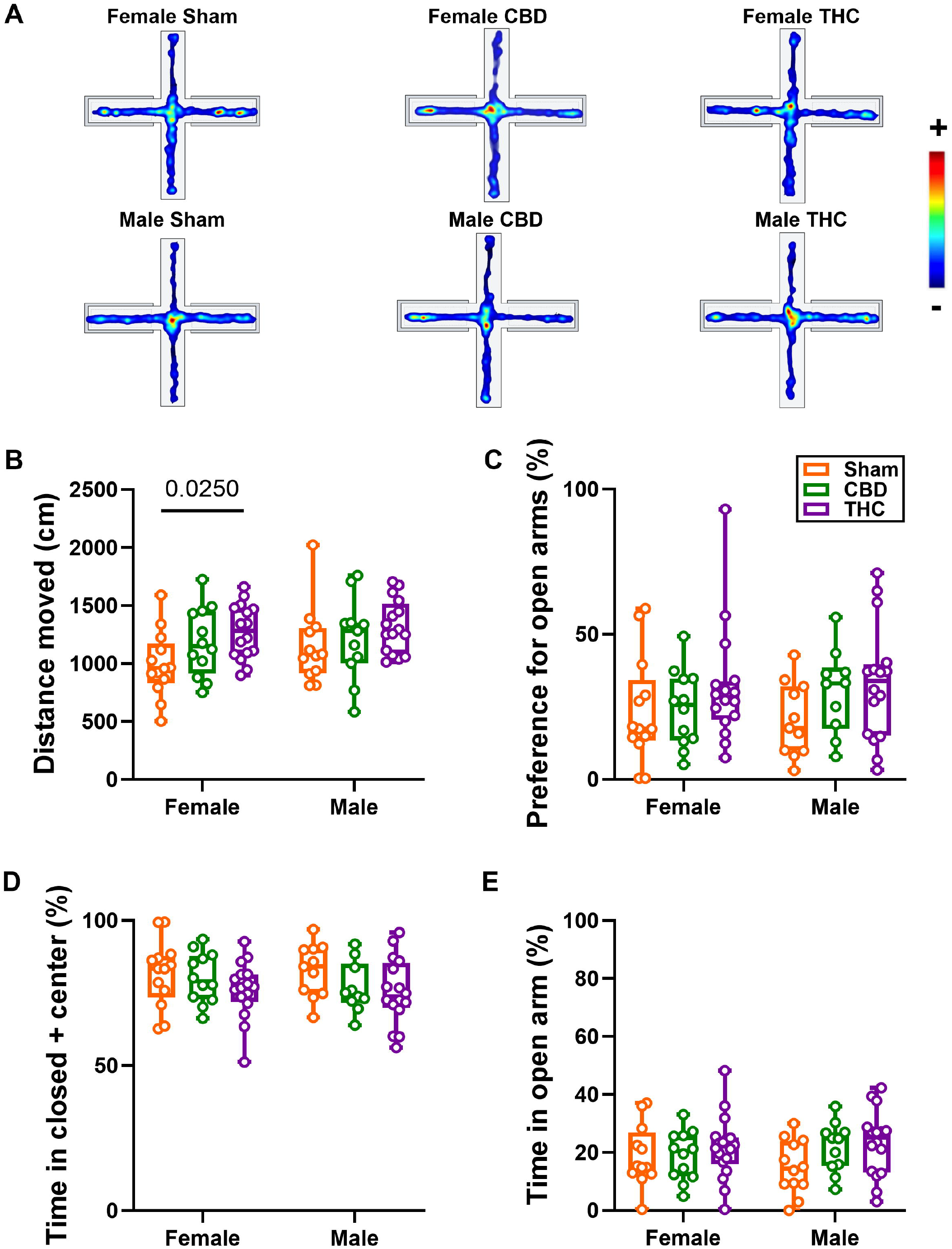
Prenatal cannabinoid (THC or CBD) exposure does not affect open-arm time in the Elevated Plus Maze in adult offspring. (A) Heatmap showing the time spent in each arm of the maze over a 300-second trial. Warm colors (e.g., red) indicate areas visited most frequently, while cool colors (e.g., blue) denote areas visited least frequently. (B–E) Quantitative analyses include total distance moved (B), preference for open arms (C), time spent in the closed arm and center (D), and time spent in the open arms (E). Notably, THC-exposed female offspring displayed increased locomotor activity, as reflected in increased distance moved (B). However, analysis of relative time spent in different maze zones revealed no significant differences between groups, regardless of sex or prenatal exposure to CBD or THC. Data in panels (C–E) are presented as box-and-whisker plots (minimum, maximum, median), with each point representing an individual animal. Statistical comparisons were performed using two-way ANOVA followed by Šídák’s multiple comparisons test. No significant effects were detected. Group sizes were as follows: Sham male (N = 12), CBD male (N = 11), THC male (N = 16), Sham female (N = 13), CBD female (N = 12), THC female (N = 16).

### Prenatal CBD, but not THC, induces enduring female-specific increases in risk-assessment postures in the EPM

To further evaluate the impact of prenatal cannabinoid exposure on exploratory strategies, we examined postural states during the elevated plus maze (Figure 2) (Iezzi, Cáceres-Rodríguez, Chavis, et al. 2025). Across groups, animals most frequently adopted a neutral stance, but females exposed to CBD displayed this posture more often than Sham controls. When focusing on stretched-attend posture (SAP), CBD-exposed females engaged in this risk-assessment behavior significantly longer than both Sham females and CBD-exposed males, an effect not reproduced in THC-exposed groups. In parallel, CBD offspring showed a reduction in contracted posture, consistent with a behavioral shift toward SAP. These findings indicate that prenatal CBD, but not THC, promotes risk-assessment behavior in adult females.

**Figure 2.**
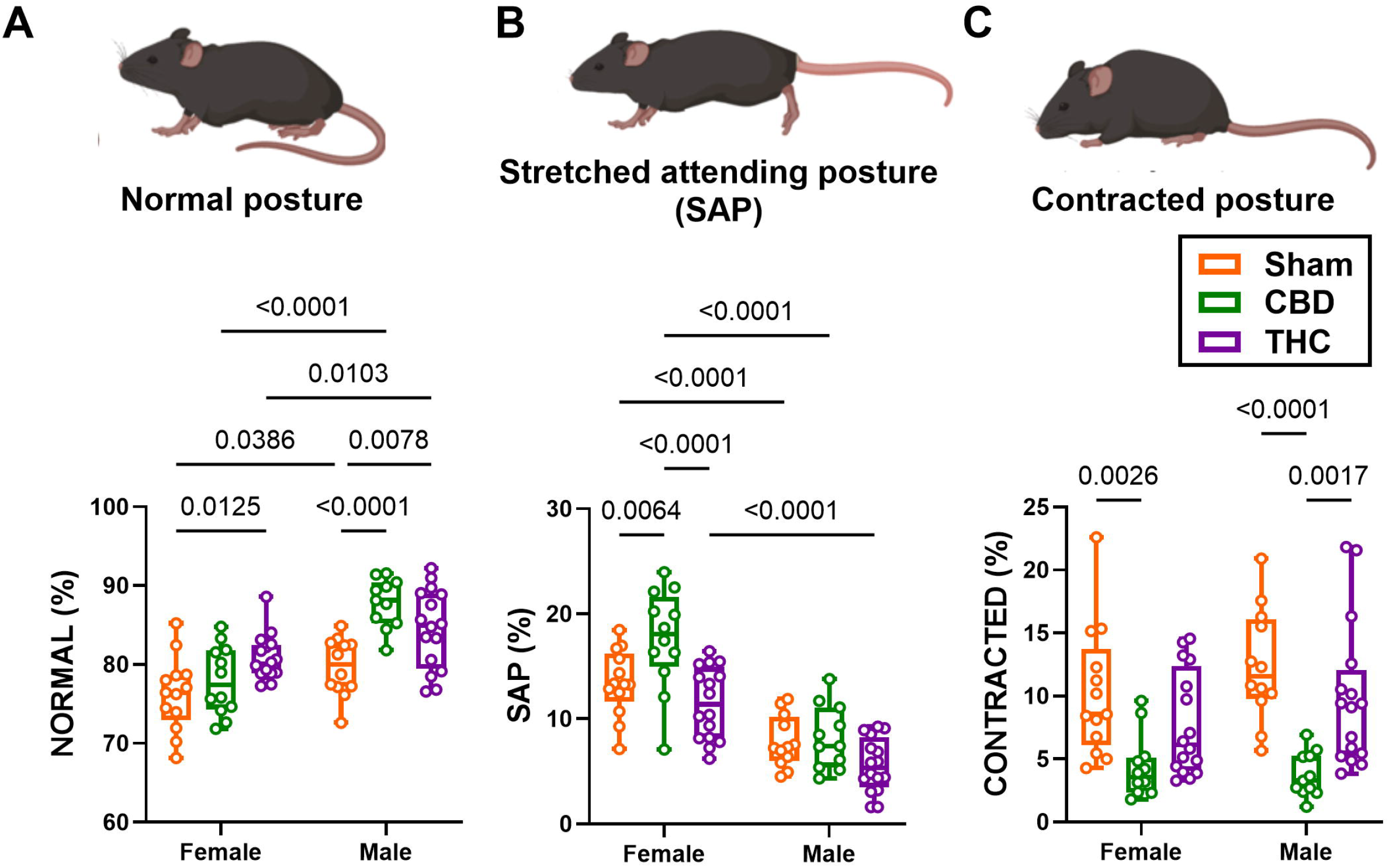
Prenatal exposure to CBD, but not THC, selectively enhances risk assessment behavior in adult female offspring. Postural analysis during the EPM task categorized behavior into three distinct states: normal posture (A), stretched-attend posture (SAP; B), and contracted posture (C). (A) Female offspring exposed to prenatal CBD exhibited a higher prevalence of normal posture. (B) Analysis revealed that CBD-exposed females spent significantly more time in SAP compared to both Sham females and CBD-exposed males — an effect absent in THC-exposed groups. (C) CBD-exposed animals also spent less time in contracted posture, consistent with an increased expression of SAP-related risk assessment behavior. Statistical comparisons were performed using two-way ANOVA followed by Šídák’s multiple comparisons test. *P values < 0.05 are indicated in the graph. Group sizes were as follows: Sham male (N = 12), CBD male (N = 11), THC male (N = 16), Sham female (N = 13), CBD female (N = 12), THC female (N = 16).

We next asked whether these posture changes were uniformly expressed across the maze or confined to specific zones (Figure 3). Independent of treatment or sex, animals spent more time in neutral postures within the closed arms compared to the open or center arms. Strikingly, CBD-exposed females displayed a selective increase in SAP when positioned in the closed arms, particularly before transitions toward the center. This spatial bias suggests that the enhancement of risk-assessment behavior following prenatal CBD exposure is most evident in contexts associated with relative safety.

**Figure 3.**
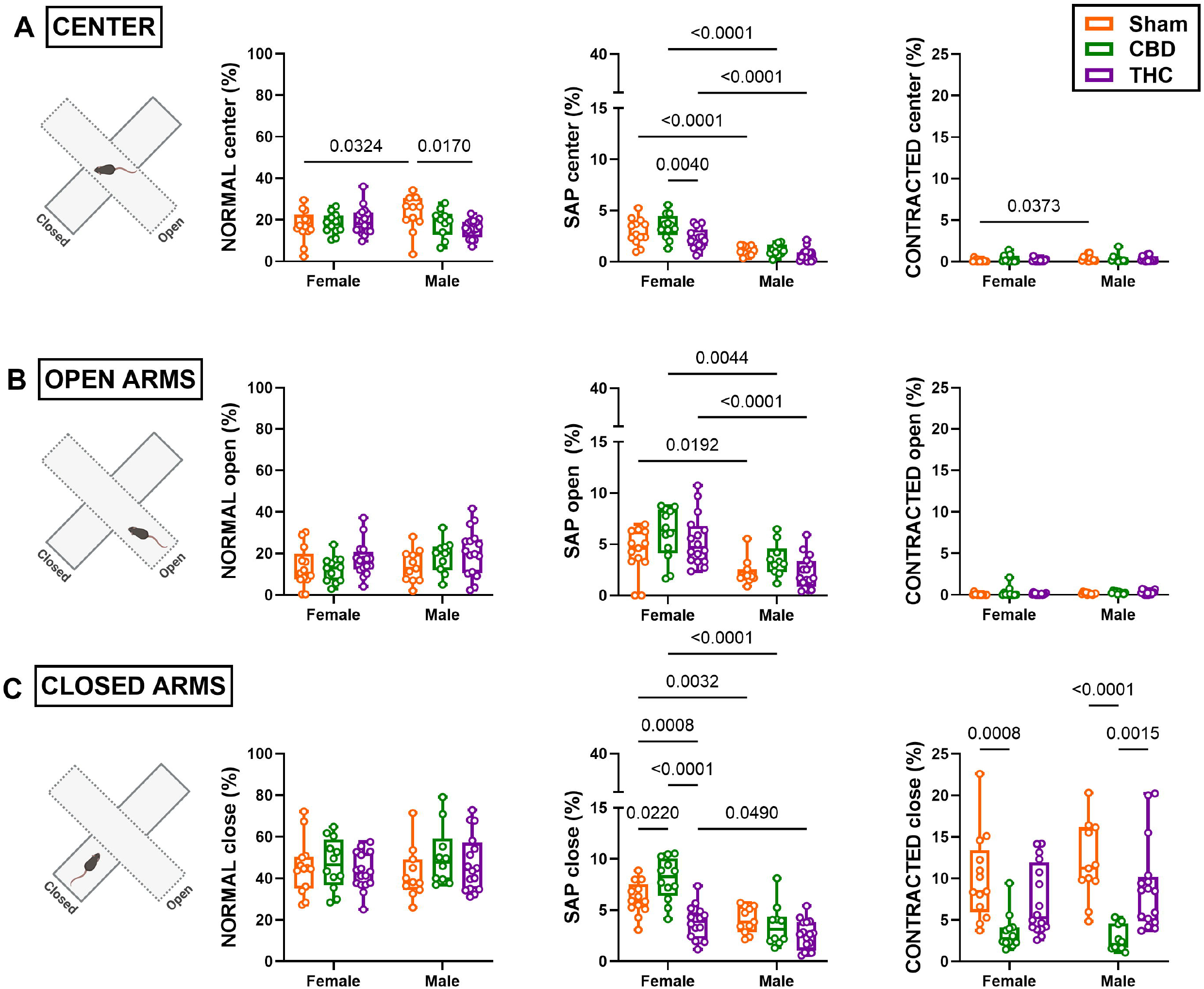
CBD-induced enhancement of risk assessment behavior in female offspring is spatially restricted to the closed arms of the maze. (A–C) Quantitative analysis of postural states—normal, contracted, and stretched-attend posture (SAP)—across the three zones of the EPM revealed that, regardless of sex or prenatal treatment, both Sham and CBD-exposed offspring spent significantly more time in normal posture within the closed arms compared to the open and center arms. Notably, CBD-exposed progeny, particularly females, exhibited elevated SAP rates specifically from the closed arms prior to center entry. Data are presented as box-and-whisker plots (minimum, maximum, median), with each point representing an individual animal. Statistical analysis was performed using two-way ANOVA followed by Šídák’s multiple comparisons test. *P values < 0.05 are indicated in the graph. Group sizes were as follows: Sham male (N = 12), CBD male (N = 11), THC male (N = 16), Sham female (N = 13), CBD female (N = 12), THC female (N = 16).

**Figure 4.**
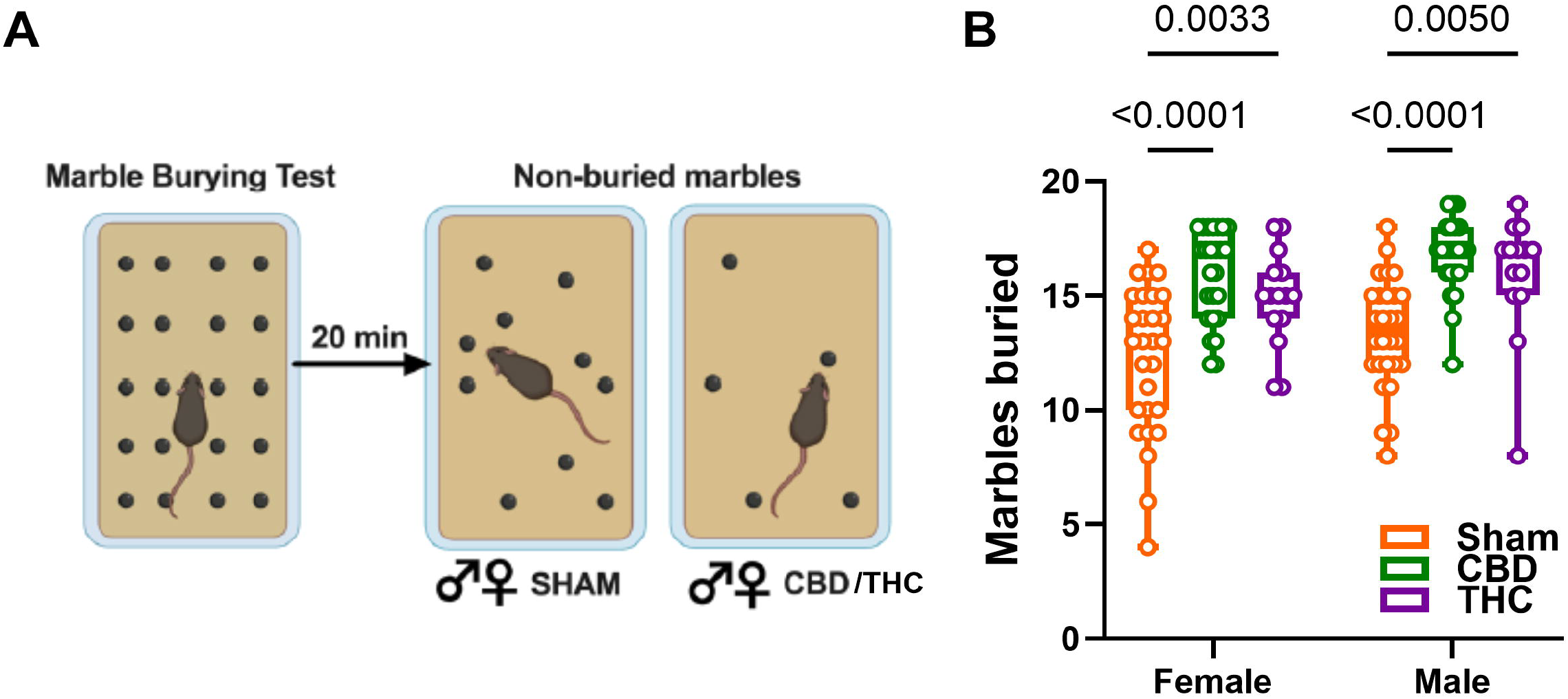
Gestational Cannabinoid Exposure Elevates Repetitive Responses in the Marble Burying Task. (A) Left: schematic representation of the marble burying paradigm. Right: representative results from Sham, CBD-, and THC-exposed animals of both sexes. (B) Quantification of buried marbles revealed that both male and female offspring prenatally exposed to CBD or THC buried significantly more marbles than their Sham counterparts. (C) Data are presented as box-and-whisker plots (minimum, maximum, median), with each point representing an individual animal. Statistical analysis was performed using two-way ANOVA followed by Šídák’s multiple comparisons test. *P values < 0.05 are indicated in the graph. Group sizes were: Sham male (N = 29), CBD male (N = 23), THC male (N = 14), Sham female (N = 28), CBD female (N = 25), THC female (N = 16).

### Prenatal CBD, but not THC, increases defensive marble burying behavior in adulthood

The MB test was used as an ethological measure of defensive behavior, in which marbles are treated as potential threats and neutralized through burying (Rodgers and Dalvi 1997; de Brouwer et al. 2019). This assay complements the EPM by probing a distinct dimension of emotional reactivity centered on repetitive defensive responding. In our previous work, we reported that prenatal CBD exposure increased marble burying in female offspring (Iezzi, Cáceres-Rodríguez, Chavis, et al. 2025), whereas the impact of prenatal THC on this behavior had not been assessed. To extend these findings, we applied the same prenatal exposure protocol to the present cohort. Analysis revealed that offspring exposed to either CBD or THC buried significantly more marbles than Sham controls. This increase was evident in both males and females. By contrast, Sham progeny of both sexes buried a comparable number of marbles, indicating no baseline sex difference in this measure.

## Discussion

Exposure to cannabis during critical developmental windows has been associated with long-term vulnerability to neuropsychiatric disorders (Navarrete et al. 2020; Scheyer et al. 2019; Bara et al. 2021; Hurd et al. 2019). In this study, we examined the consequences of prenatal cannabinoid exposure on adult behavior using two complementary paradigms: the EPM and the MB test. Together, these assays allowed us to probe both anxiety-related approach–avoidance behavior and defensive/repetitive responding.

Our findings reveal a nuanced picture. Neither prenatal CBD nor THC exposure altered the classical EPM measure of anxiety—time spent in the open arms. This is consistent with our previous report in rats showing that prenatal THC did not affect adult open-arm exploration (Bara et al., 2018). These convergent results suggest that gross indices of approach–avoidance conflict may not be sensitive to the long-term consequences of gestational cannabinoid exposure.

However, ethological analysis revealed a more specific pattern: prenatal CBD selectively increased SAP, a well-established risk-assessment behavior linked to heightened vigilance and anxiety in rodents (Holly et al. 2016; Iezzi, Cáceres-Rodríguez, Chavis, et al. 2025). This effect was restricted to adult female offspring and reflects a change in the microstructure of risk assessment rather than a generalized increase in anxiety. CBD-exposed females spent more time in SAP and less time in contracted posture, with the SAP enhancement most pronounced when mice initiated movement from the closed arms toward the centre. No comparable SAP effect was observed in THC-exposed groups, indicating a compound-specific, female-biased developmental liability.

By contrast, the marble-burying MB, revealed a robust, sex-independent increase in repetitive defensive responding following prenatal exposure to either CBD or THC. Offspring of both sexes buried more marbles than Sham controls, indicating that repetitive or compulsive-like defensive strategies (Witkin 2008; de Brouwer et al. 2019; Rodgers and Dalvi 1997) are broadly sensitive to gestational cannabinoid perturbation. The parallel MB effect for CBD and THC, contrasted with the female-restricted increase in SAP, implies at least two separable behavioral outcomes of prenatal cannabinoid exposure.

Taken together, these results highlight two distinct trajectories of vulnerability: (i) a sex-specific, CBD-driven increase in risk-assessment behaviors that may model heightened anxiety vigilance in females, and (ii) a general increase in defensive/repetitive behaviors following both CBD and THC exposure. This dual pattern aligns with clinical evidence that anxiety and compulsivity often co-occur but can also diverge in their developmental origins (Göbel et al. 2022; Havewala et al. 2022).

These empirical patterns are compatible with multiple, non-exclusive mechanistic routes. Prenatal engagement of cannabinoid signalling can perturb axon guidance, synaptogenesis, and synaptic plasticity within prefrontal–limbic and corticostriatal networks (Scheyer et al. 2019); such circuit-level rewiring is likely to alter fine-grained defensive tactics (for example, SAP) as well as action-selection processes that support repetitive responding. Sex differences may arise from interactions between endocannabinoid signalling and sex-dependent modulators of plasticity or hormonal trajectories that preferentially shape vigilance-related behaviors in females (Aspesi et al. 2025). Maternal–fetal influences, including changes in placental function, maternal stress responsivity, or maternal care, could also contribute to the observed outcomes (Simone et al. 2022; Bernabeu et al. 2023).

The dissociation between posture-based and coarse EPM readouts highlights the value of ethological, multidimensional behavioral batteries: posture metrics can uncover subtle, sex-specific changes that escape conventional measures such as open-arm time.

Limitations of the present study include reliance on a single exposure regimen—chosen deliberately to represent a low dose—and a restricted behavioral panel. Dose–response relationships, the timing of exposure across gestation, and the generalizability of these findings to other tasks and species remain to be determined. Future studies should incorporate physiological readouts (HPA-axis function, autonomic measures), broaden behavioral phenotyping, and directly relate ethological phenotypes to cellular and circuit markers (neuronal morphology and intrinsic properties, synaptic physiology, receptor expression), with explicit attention to sex differences.

In sum, prenatal exposure to major cannabinoids does not produce a uniform increase in classical anxiety readouts but yields compound- and sex-specific alterations in defensive behavior. Prenatal CBD selectively altered the microstructure of risk assessment in adult females, indexed by increased SAP and reduced contracted posture in the EPM, whereas both CBD and THC produced a sex-independent elevation of repetitive defensive responding in the MB assay. From a translational standpoint, these results caution against assuming prenatal CBD is benign; although CBD and THC differ in the specificity of their effects, both compounds produced enduring changes in defensive and repetitive behavioral strategies in adult offspring.

## Author Contributions

A.C.-R.: conceptualization, data curation, formal analysis, validation, writing—review and editing. D.I.: data curation, writing—review and editing. P.C.: conceptualization, supervision. O.J.J.M.: conceptualization, supervision, funding acquisition, methodology, project administration, writing—original draft, review, and editing. All authors have read and agreed to the published version of the manuscript.

## Acknowledgements

The authors are grateful to the Chavis-Manzoni team members for helpful discussions.

