## Supplementary material for "Prenatal Cannabinoids Produce Sex-Specific Changes in Risk Assessment and Shared Increases in Repetitive Behavior in Adult Offspring": Table 1

| **Measure** | **Treatment** | **Median** | **Max** | **Min** | **N** | ***Two-way ANOVA Main effects and interactions*** | ***Multiple comparison (Šidák’s post hoc test)*** |
| --- | --- | --- | --- | --- | --- | --- | --- |
| **Distance moved (cm)** | SHAM FEMALE | 957.618 | 1591.55 | 503.066 | 13 | **Interaction:** F (2, 74) = 0,3939; P=0,6758 **Sex:** F (1, 74) = 1,398; P=0,2409 **Treatment** F (2, 74) = 4,419; P=0,0154 | FEMALE "Sham vs. CBD" P=0,2099 "Sham vs. THC" P=0,0250 "CBD vs. THC" P=0,6904 MALE "Sham vs. CBD" P=0,8485 "Sham vs. THC" P=0,2752 "CBD vs. THC" P=0,6260 |
|  | CBD FEMALE | 1150.015 | 1723.54 | 750.624 | 12 |  |  |
|  | THC FEMALE | 1282.48 | 1661.58 | 899.255 | 16 |  |  |
|  | SHAM MALE | 1075.535 | 2020.9 | 810.543 | 11 |  |  |
|  | CBD MALE | 1282.85 | 1758.49 | 584.972 | 10 |  |  |
|  | THC MALE | 1276.36 | 1705.87 | 1012.67 | 16 |  |  |
| **Preference for Open Arms (%)** | SHAM FEMALE | 17.39257 | 58.8596 | 0.383216 | 13 | **Interaction:** F (2, 72) = 0,3889; P=0,6792 **Sex:** F (1, 72) = 0,1030; P=0,7492 **Treatment** F (2, 72) = 2,464; P=0,0922 | FEMALE "Sham vs. CBD" P=0,9909 "Sham vs. THC" P=0,4119 "CBD vs. THC" P=0,5080 MALE "Sham vs. CBD" P=0,3921 "Sham vs. THC" P=0,1629 "CBD vs. THC" P=0,9285 |
|  | CBD FEMALE | 25.64002 | 49.27898 | 5.156046 | 12 |  |  |
|  | THC FEMALE | 28.300745 | 93.13346 | 7.419347 | 16 |  |  |
|  | SHAM MALE | 17.72665 | 42.86245 | 3.019399 | 11 |  |  |
|  | CBD MALE | 33.124375 | 55.86683 | 7.895273 | 10 |  |  |
|  | THC MALE | 33.847205 | 71.15472 | 3.166812 | 16 |  |  |
| **Closed Arms + Centre (%)** | SHAM FEMALE | 83.4482 | 99.41843 | 62.6866 | 13 | **Interaction:** F (2, 72) = 0,2805; P=0,7563 **Sex:** F (1, 72) = 0,2130; P=0,9357 **Treatment** F (2, 72) = 2,798; P=0,0676 | FEMALE "Sham vs. CBD" P=0,9243 "Sham vs. THC" P=0,3009 "CBD vs. THC" P=0,5406 MALE "Sham vs. CBD" P=0,3705 "Sham vs. THC" P=0,1621 "CBD vs. THC" P=0,9430 |
|  | CBD FEMALE | 79.02395 | 93.5023 | 66.329 | 12 |  |  |
|  | THC FEMALE | 76.99995 | 92.7134 | 51.2264 | 16 |  |  |
|  | SHAM MALE | 84.0974 | 96.83627 | 66.5836 | 11 |  |  |
|  | CBD MALE | 74.4891 | 91.90925 | 63.9245 | 10 |  |  |
|  | THC MALE | 73.7754 | 95.84524 | 56.1791 | 16 |  |  |
| **Open Arms (%)** | SHAM FEMALE | 15.09575 | 37.0318 | 0.387722 | 13 | **Interaction:** F (2, 75) = 0,8071; P=0,4500 **Sex:** F (1, 75) = 0,006545; P=0,6458 **Treatment** F (2, 75) = 2,087; P=0,1312 | FEMALE "Sham vs. CBD" P=0,9997 "Sham vs. THC" P=0,7523 "CBD vs. THC" P=0,7379 MALE "Sham vs. CBD" P=0,9997 "Sham vs. THC" P=0,7523 "CBD vs. THC" P=0,7379 |
|  | CBD FEMALE | 20.3919 | 33.0084 | 4.90141 | 12 |  |  |
|  | THC FEMALE | 22.04415 | 48.213 | 0.381558 | 16 |  |  |
|  | SHAM MALE | 14.3997 | 29.9907 | 0 | 11 |  |  |
|  | CBD MALE | 24.776 | 35.8404 | 7.31569 | 10 |  |  |
|  | THC MALE | 25.2479 | 42.2859 | 3.07094 | 16 |  |  |
